# DEAH-box helicase 37 (*DHX37*) defects are a novel molecular etiology of 46,XY gonadal dysgenesis spectrum

**DOI:** 10.1101/477992

**Authors:** Thatiana E. da Silva, Nathalia L. Gomes, Antonio M. Lerario, Catherine E. Keegan, Mirian Y. Nishi, Filomena M. Carvalho, Eric Vilain, Hayk Barseghyan, Alejandro Martinez-Aguayo, María V. Forclaz, Regina Papazian, Luciani R. Carvalho, Elaine M. F. Costa, Berenice B. Mendonca, Sorahia Domenice

## Abstract

46,XY gonadal dysgenesis is a heterogeneous disorder of sex development (DSD) that features abnormal gonadal development and varying degrees of undervirilization of the external genitalia, ranging from micropenis to female-like genitalia. Embryonic testicular regression syndrome (ETRS; MIM: 273250) is considered part of the clinical spectrum of 46,XY gonadal dysgenesis. Most ETRS patients present micropenis or atypical genitalia associated with a complete absence of gonadal tissue in one or both sides. In most patients with gonadal dysgenesis, the genetic diagnosis is unclear. We performed whole exome sequencing in ETRS patients and identified a rare variant, the p.Arg308Gln, in DEAH (Asp-Glu-Ala-His) box polypeptide 37 (*DHX37*) in 5 affected individuals from three unrelated families. We expanded the analysis of *DHX37* coding region to additional 71 patients with 46,XY gonadal dysgenesis and identified the p.Arg308Gln and three other *DHX37* missense variants (p.Arg151Trp, p.Thr304Met and p.Arg674Trp) in 11 affected members from eight distinct families (8 patients with ETRS, two with partial gonadal dysgenesis and one 46,XY DSD female patient previously gonadectomized). The p.Arg308Gln and p.Arg674Trp recurrent variants were identified in six and three families, respectively. Segregation analysis revealed sex-limited autosomal dominant inheritance in 4 families, autosomal dominant with incomplete penetrance in one family and autosomal recessive in another family. Immunohistochemical analysis of normal testes revealed that DHX37 is expressed in germ cells at different stages of maturation.

This study demonstrates an expressive frequency of rare predicted to be deleterious *DHX37* variants in 46,XY gonadal dysgenesis group, particularly those individuals exhibiting the ETRS phenotype (25% and 50%, respectively).

Our findings indicate that *DHX37* is a new player in the complex cascade of male gonadal differentiation and maintenance, thus establishing a novel and frequent molecular etiology for 46,XY gonadal dysgenesis spectrum, mainly for embryonic testicular regression syndrome.

## Introduction

46, XY gonadal dysgenesis is a heterogeneous disorder of sex development (DSD) that features abnormal gonadal development and varying degrees of undervirilization of the external genitalia, ranging from micropenis to female-like genitalia. The gonads from these patients present a wide array/spectrum of histological abnormalities, ranging from disorganized seminiferous tubules and the presence of an ovarian-like stroma, to complete absence of testicular tissue. Müllerian structures may be partially or fully developed in these patients^1^. Embryonic testicular regression syndrome (ETRS; MIM: 273250) is considered part of the clinical spectrum of 46,XY gonadal dysgenesis. Most ETRS patients present micropenis or atypical genitalia associated with a complete absence of gonadal tissue in one or both sides. The presence of micropenis suggests that functioning testes were present during early fetal development, but disappeared before birth^2,3^.

Several genetic defects have been described in sporadic and familial forms of 46,XY gonadal dysgenesis, but the genetic etiology of most patients remains unknown^4–8^.

Here, we used next generation-sequencing based methods, including whole-exome sequencing and targeted gene panels, to investigate the underlying genetic etiology in a large cohort of 46,XY patients with gonadal dysgenesis of unknown cause. We identified recurrent rare variants in DEAH (Asp-Glu-Ala-His) box polypeptide 37 (*DHX37*) in several affected individuals from distinct families, establishing a novel genetic cause for 46,XY gonadal dysgenesis spectrum.

### Ethics

This study was approved by the Ethics Committee of the Hospital das Clínicas da Faculdade de Medicina da Universidade de São Paulo, the Institutional Review Board of the University of Michigan Medical School, the Hospital de Garrahan Escuela de Medicina, Pontificia Universidad Católica de Chile, and the Hospital Nacional Prof. Dr. A. Posadas, Buenos Aires, Argentina. Written informed consent was obtained from all patients, their parents or legal guardians.

### Subjects and Methods

Four children from two Brazilian families (Family 1 and 2) and one Chinese-American child with ETRS were firstly studied by whole-exome sequencing. Two additional families with ETRS (one from Chile, the other from Argentina) (Table 1 and 2) and 71 Brazilian patients with sporadic 46,XY DSD (39 patients with partial and complete 46,XY gonadal dysgenesis (atypical genitalia, presence of Mullerian derivatives and/or at least one dysgenetic gonad or absence of gonadal tissue) and thirty-two 46,XY DSD patients with unknown etiology (molecular defects in LHCG and androgen receptor genes and *5ARD2* gene were ruled out by Sanger method as well as testosterone synthesis defects by hormonal evaluation) or due to previously gonadectomy. All patients have a normal GTG-banded metaphases 46,XY karyotype.

**Table 1.** Phenotype of familial 46,XY DSD patients with gonadal dysgenesis associated with heterozygous likely pathogenic *DHX37* variants

| Origin<br>Parameters |  | Brazilian |  | Brazilian |  | Chilean |  | Argentinian |  |  |
| --- | --- | --- | --- | --- | --- | --- | --- | --- | --- | --- |
| Subject |  | F1:III-2 | F1:III-3 | F2:II-4 | F2:III-1 | F3:II-1 | F3:II-2 | F4:III-1 | F4:III-2 | F4:II-4 |
| Sex of rearing |  | Male | Male | Male | Female | Male | Male | Male | Male | Male |
| Age at presentation (yrs) |  | 2.2 | 1.8 | 14.0 | 1.8 | 0.6 | 0.08 | Newborn | 4 | 2 |
| Diagnosis |  | ETRS | ETRS | ETRS | ETRS | ETRS | ETRS | ETRS | ETRS | PGD |
| External genitalia |  | Micropenis | Micropenis | Micropenis | Micropenis | Micropenis | Micropenis | Micropenis | Micropenis | Micropenis |
| Gonads |  | Non-palpable | Non-palpable | Non-palpable | Non-palpable | Non-palpable | Non-palpable | Non-palpable | Non-palpable | Left testis – scrotum |
| Histologic analysis |  | Bilateral dysgenetic gonads | L: no gonadal tissue<br>R: dysgenetic gonad | No gonadal tissue | L: no gonadal tissue<br>R: dysgenetic gonad | Bilateral dysgenetic gonads | Bilateral dysgenetic gonads | No gonadal tissue | No gonadal tissue | R: no gonadal tissue<br>L: dysgenetic gonad with IUGN* |
| Wolff derivatives |  | Present | Present | Present | Present | Present | Present | Present | Present | Present |
| Mullerian Derivatives | Tubes | Present | Present | Present | Present | Absent | Present | Present | Present | Absent |
|  | Uterus | Absent | Absent | Absent | Absent | Absent | Absent | Absent | Absent | Absent |
| LH (U/L) |  | 14.5 | 12 | 3.5 | 1.9 | 0.25 | 0.17 | <0.5 | <0.5 | 26 |
| FSH (U/L) |  | 117 | 133 | 87 | 56 | 10.9 | 9.5 | NA | 9 | 112 |
| Basal T (ng/dL) |  | <10 | <10 | <10 | NA | 16 | <10 | 38 | 27 | 16 |
| T after hCG test (ng/dL) |  | <10 | NA | 29 | 26 | 18 | <10 | 40 | 29 | NA |
| Allelic variants |  | p.Arg308Gln | p.Arg308Gln | p.Arg308Gln | p.Arg308Gln | p.Arg674Trp | p.Arg674Trp | p.Arg674Trp | p.Arg674Trp | p.Arg674Trp |
| Variants state |  | Heterozygous | Heterozygous | Heterozygous | Heterozygous | Heterozygous | Heterozygous | Heterozygous | Heterozygous | Heterozygous |

NA- not available; PGD- partial gonadal dysgenesis; IUGN- intratubular undifferentiated germinal neoplasia; \* testicular biopsy
L: left; R: right; Conversion factors to SI units: T: testosterone; T ng/dL to nmol/L, multiply by 0.0347

**Table 2.** Phenotype of 46,XY DSD patients with sporadic gonadal dysgenesis and likely pathogenic *DHX37* allelic variants.

| Origin<br>Parameters |  | Chinese-American | Brazilian |  |  |  |  |  |
| --- | --- | --- | --- | --- | --- | --- | --- | --- |
| Patient |  | F5:II-1 | F6:II-1 | F7:II-1 | F8:II-1 | F9:II-5 | F10:II-1 | F11:II-1 |
| Social sex |  | Male | Male to Female | Male to Female | Female | Female | Male to female | Female |
| Age at presentation (yrs) |  | 0.18 | 30 | 4.3 | 7.7 | 35 | 19 | 3.7 |
| Diagnosis |  | ETRS | ETRS | ETRS | PGD | Previous gonadectomy | ETRS | PGD |
| External genitalia |  | Micropenis | Micropenis | Micropenis | Female | Previous genitoplasty | Micropenis | Atypical |
| Gonads |  | Non-palpable | Non-palpable | Non-palpable | Non-palpable | NA | Non-palpable | Non-palpable |
| Histological analysis |  | No gonadal tissue | No gonadal tissue | R: no gonadal tissue<br>L: dysgenetic gonad | Bilateral dysgenetic gonads | NA | No gonadal tissue | Bilateral dysgenetic gonads |
| Wolff derivatives |  | Present | Present | Present | Present | NA | Present | Present |
| Mullerian Derivatives | Tubes | Absent | Rudimentary | Present | Absent | NA | Absent | Rudimentary |
|  | Uterus | Absent | Absent | Absent | Absent | Absent | Absent | Absent |
| LH (U/L) |  | 0.1 | 10 | 1 | 0.1 | NA | 23 | 0.1 |
| FSH (U/L) |  | 0.4 | 40 | 48.5 | 4.89 | NA | 62 | 0.4 |
| Basal T (ng/dL) |  | <10 | NA | NA | <10 | NA | 21 | <10 |
| T after hCG test (ng/dL) |  | <10 | <10 | <10 | NA | NA | NA | 337 |
| Allelic variants |  | p.Arg308Gln | p.Arg308Gln | p.Arg151Trp* | p.Arg308Gln | p.Thr304Met | p.Arg674Trp | p.Arg674Trp |
| Variants state |  | Heterozygous | Heterozygous | Homozygous | Heterozygous | Heterozygous | Heterozygous | Heterozygous |

NA: not available; PGD- Partial gonadal dysgenesis; Conversion factors to SI units: T, ng/dL to nmol/L, multiply by 0.0347.
\*Variant of uncertain significance

### Molecular Analysis

Molecular studies were performed by targeted massively parallel sequencing or Sanger sequencing. Details of DNA extraction, Sanger and massive parallel sequencing, variant filtering and immunohistochemical staining analysis are available in the Supplementary Data.

Molecular analysis was performed on genomic DNA obtained from peripheral leukocytes. The identified allelic variants in massive parallel sequencing were subsequently confirmed by Sanger method as well as segregation analysis of the candidate variants. Allelic variants were considered to be deleterious when they were predicted to be pathogenic by at least three of the following *in silico* algorithms (SIFT, PolyPhen2, Mutation Taster, Mutation Assessor, FATHMM, LRT, LR score, and radial SVM) and by the conservation scores (GERP++, SiPhy, PhyloP, CADD score). In order to select disease-causing variants, we excluded those with a minor allele frequency (> 0.5%) in publicly available population databases (gnomAD, 1000 Genomes, dbSNP, ExAC, ESP6500, ABraOM-Brazilian Genomic Variants) and in 548 Brazilian in-house controls.

Protein interaction networks (National Center for Biotechnology Information [NCBI], Biogrid, and String databases) were used to evaluate interaction among proteins which are co-expressed with DHX37 in testicular cells. Using protein interactions we constructed an hypothetical cascade of testicular differentiation including DHX37.

### Histological analysis

One single pathologist (FMC) reviewed the histology of the gonadal tissue samples from all Brazilian patients.

### Immunohistochemical staining study

Eight formalin-fixed paraffin-embedded testicular necropsy samples from 46,XY individuals with different chronological ages (27 and 33 weeks gestational age, 1, 53, and 180 days of age, 13, 23 and 53 years of age) were collected and used for DHX37 expression analysis by immunohistochemistry.

### Statistical analysis

To provide additional genetic evidence of the association of *DHX37* with gonadal dysgenesis phenotype, we performed aggregate variant analyses comparing allele frequencies between our cohort of 46,XY DSD patients and public databases (gnomAD and ABraOM). In public databases, we selected variants with similar characteristics of the *DHX37* variants observed in our cohort: rare nonsynonymous variants (minor allele frequency of 0.01) located in the two highly conserved protein domains (ATP-binding and Helicase C-terminal domains) that are predicted to be pathogenic by at least four *in silic*o tools (PROVEAN, SIFT, PolyPhen-2, and Mutation Assessor). Allele frequency differences between groups were analyzed by x2 test, and statistical significance was set at P<0.05. Statistical analyses were performed by using the SIGMAstat statistical software package (Windows version 4.0; SPSS Inc., San Rafael, CA).

## Results

### Exome sequencing nominated DHX37 as a candidate gene in three unrelated families

The exome sequencing data identified the single missense *DHX37* variant, c.923C>T [p.Arg308Gln]; GenBank: NM_032656.3, in three unrelated families with 46,XY gonadal dysgenesis (Figure 1). *DHX37 is* a member of the large DEAH family of proteins and encodes an RNA helicase^11,12^. The DHX37 protein is comprised by 1157 amino acids and four main domains, including the conserved motifs of the helicase core region; the Helicase ATP-binding (position 262-429) and the Helicase superfamily c-terminal domain (position 585-674), in addition to the helicase associated domain (position 768-859) and the oligonucleotide/oligosaccharide-binding-fold (position 894-1011) (Figure 2). The p.Arg308Gln variant is predicted to be pathogenic by six *in-silico* prediction sites (Table S2) and is not present in genomic population databases, except in the gnomAD database, where this variant was found at a very low allele frequency (0.00003) (Table S3-S4). The p.Arg308Gln variant was not identified in Brazilian population: in 548 individuals by Sanger sequencing, in 308 by in-house exome and in 609 healthy Brazilian subjects from ABraOm database. Finally, our phasing analysis ruled out a founder effect in Brazilian families 1 and 2. To identify other possible pathogenic variants on *DHX37*, we further extended our study to include 81 patients with variable phenotypes of 46,XY gonadal dysgenesis. We studied the entire coding sequence of *DHX37* either by massive parallel sequencing (WES or DSD targeted panel) or Sanger sequencing (Table S1). Three other novel variants, the c.451G>A (p.Arg151Trp), the c.911C>T (p.Thr304Met) and the c.2020G>A (p.Arg674Trp) were identified in four distinct families: (Table 1 and 2, Figure 1). All were in heterozygous state, except the p.Arg151Trp variant.

**Figure 1.**
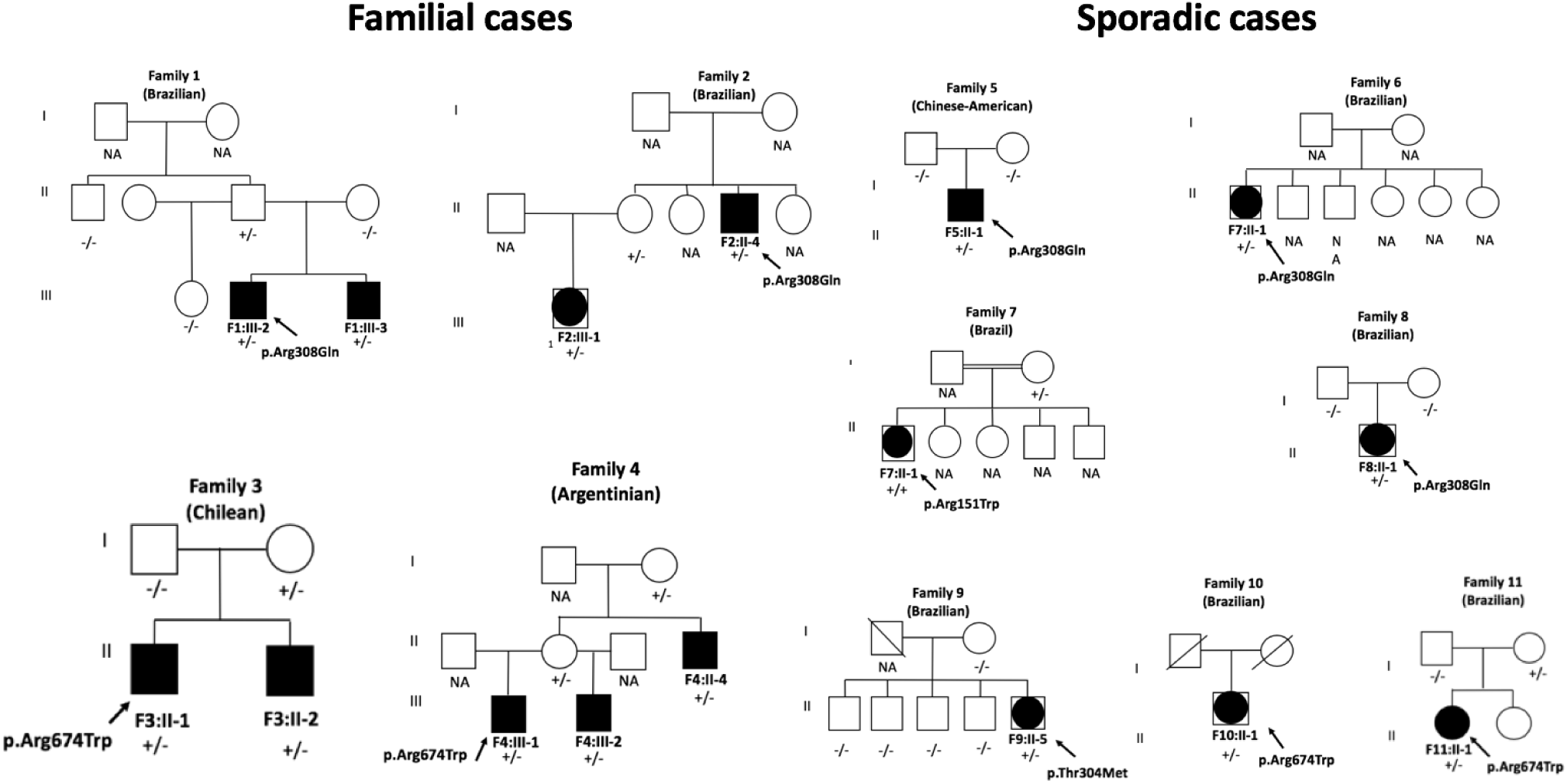
Pedigrees of the eleven families with potential disease-causing *DHX37* variants. Filled symbols represent affected individuals. The affected raised males (46,XY) are indicated by filled squares and the affected raised females (46,XY) are shown by large dark dots within the squares. Symbols with a diagonal line represent deceased individuals. The *DHX37* genotype is shown for the individuals whose DNA sample was available; +/- indicates a heterozygous state; +/+ indicates a homozygous state for the variant, and -/- indicates a homozygous state for wild-type allele. NA- DNA not available. The paternity of patient F5:II-1 and F8:II-1 was confirmed.

The homozygous p.Arg151Trp was identified in one sporadic ETRS patient. The heterozygous p.Arg308Gln variant was identified in three Brazilian patients with sporadic ETRS. The p.Arg674Trp variant was identified in the two Chilean brothers with ETRS and in the three affected members of one Argentinian family (two brothers with ETRS, their uncle with partial GD) and in one Brazilian sporadic partial GD patient. The heterozygous p.Thr304Met was found in a Brazilian patient, who underwent previous gonadectomy (Tables 1 and 2, Figure 1).

The p.Thr304Met was located inside DHX37 Helicase ATP-binding (position 262-429) and was classified as deleterious by six in-silico prediction sites (Figure 2, Table S2). The p.Thr304Met was not found in all searched population databases (Figure 2; Tables S3-S4). The p.Arg674Trp variant was located in the Motif VI (position 667-674), inside the Helicase superfamily c-terminal domain, and was classified as deleterious by six *in silico* prediction tools (Table S2) and was not identified in all searched population databases (Figure 2; Table S3-S4). The p.Arg151Trp (dbSNP: rs577400960) variant was located at the beginning of the protein sequence, outside the Helicase ATP-binding, and was classified as possibly deleterious by four *in silico* prediction tools (Figure 2; Table S2). It was found in heterozygous state in the 1000 Genome and in Brazilian controls with allele frequencies of 0.02% (1/5007). It was not identified in the ABraOM-Brazilian Genomic Variants and Brazilian in-house controls (Table S3 -S4).

**Figure 2.**
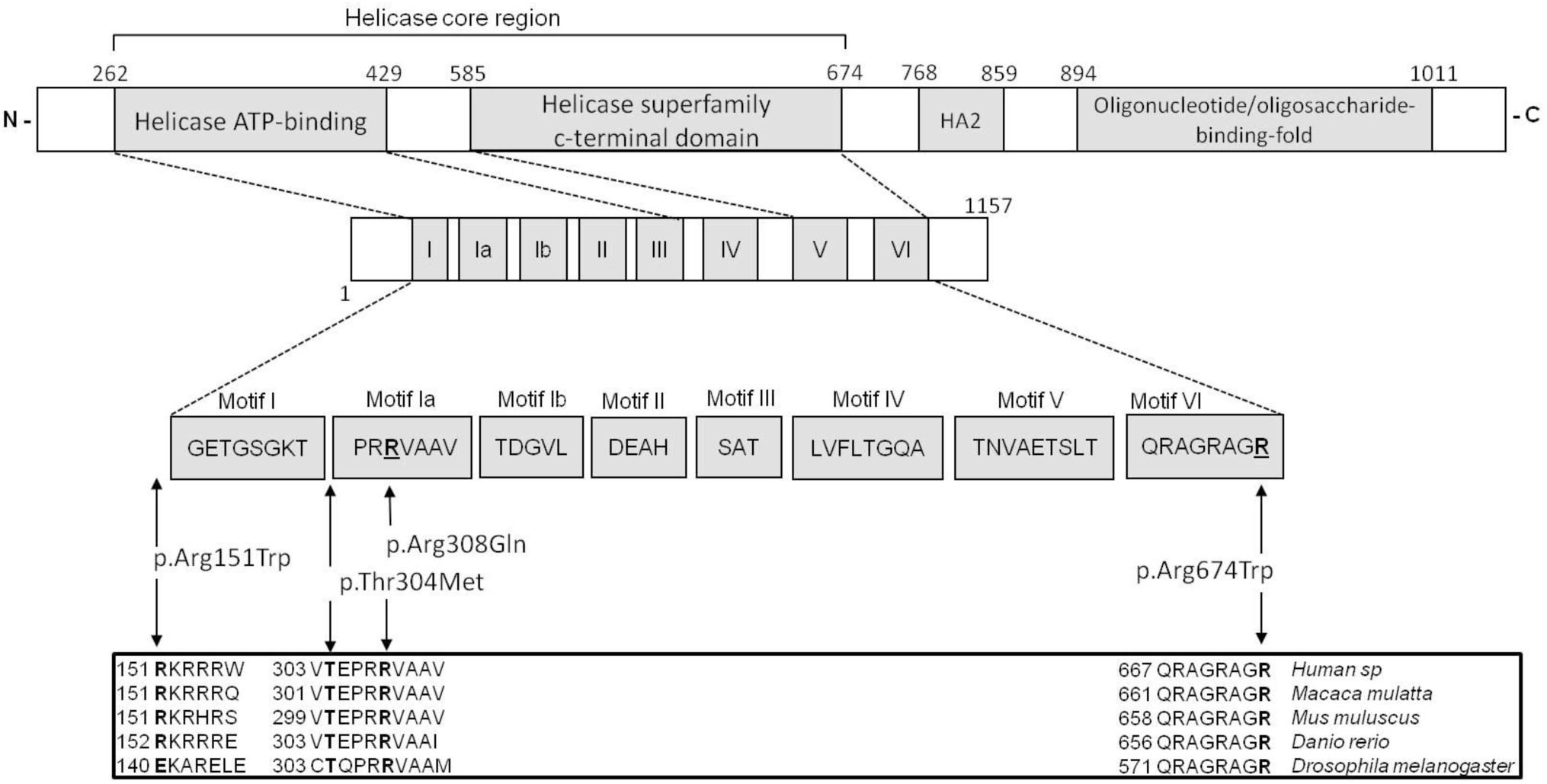
Identified variants are localized within conserved helicase domains of DHX37. Top: Schematic protein structure of DHX37 showing conserved motifs of the helicase core region, the helicase associated domain (HA2) and the oligonucleotide/oligosaccharide-binding-fold. Middle: Nucleotide-interacting motifs (I, II, and VI), nucleic acid-binding motifs (Ia, Ib, and IV), motif V, which binds nucleic acid and interacts with nucleotides, and motif III, which couples ATP hydrolysis to RNA unwinding (N-N terminus; C-C terminus). Bottom: Amino acids within conserved motifs of the helicase core region. The position of the first and last amino acid within each motif is denoted below left and right, respectively. The position of the allelic variants identified in this study are indicated with vertical arrows and shown in bold in the different species sequence, which the pathogenic variants (p.Arg308Gln, p.Thar304Met and p.Arg674Trp) are located at the Motif Ia and Motif VI.

### Frequency of the *DHX37* variants in our 46,XY DSD cohort

The allele frequency of rare *DHX37* variants predicted to be deleterious found in our whole cohort of 46,XY DSD patients (10%) was expressively higher than that observed in individuals from gnomAD (568/141456 individuals [0.004%; *p*<0.001]) and from the Brazilian cohort (1/609 individuals [0.002%; *p*<0.001]). Our study demonstrated an expressive frequency of rare predicted to be deleterious *DHX37* variants in 46,XY gonadal dysgenesis group, particularly those individuals exhibiting the ETRS phenotype (25% and 50%, respectively).

### Segregation analysis revealed different inheritance patterns

Segregation analysis of the DHX37 variants in seven families displayed a sex-limited autosomal dominant pattern in three families (F2, F3, F4). In the Family 1, the presence of the p.Arg308Gln variant in the asymptomatic father suggests an autosomal dominant pattern of inheritance with incomplete penetrance. An autosomal recessive inheritance was identified in one consanguineous family (F7) (Figure 1). In two sporadic cases (F5:II-1 and F7:II-1) the confirmed paternity displayed a *de novo* status of the p.Arg308Gln *DHX37* variant (Figure 1).

### Absence of founder effect of *DHX37* p.Arg308Gln variant

The comparative analysis of polymorphic markers located around the p.Arg308Gln variant in families 1 and 2 evaluated in exome data excluded the occurrence of founder effect.

### DHX37 protein identified in different stages of germ cells maturation

To gain further insights on the role of DHX37 in testicular development, we studied its expression in testes from newborn children and from adults by immunohistochemistry technique. DHX37 expression analysis in newborns was found exclusively in spermatogonia and it was characterized by a regular perinuclear halo pattern. The same pattern is also observed in spermatogonia of adult testis. A progressive condensation of protein around the nucleus was observed as cells changed from spermatocyte stage 1 to 2, generating a localized paranuclear pattern. A punctate perinuclear pattern of expression was observed in spermatids, but no staining was observed in spermatozoa (Figure 3).

**Figure 3.**
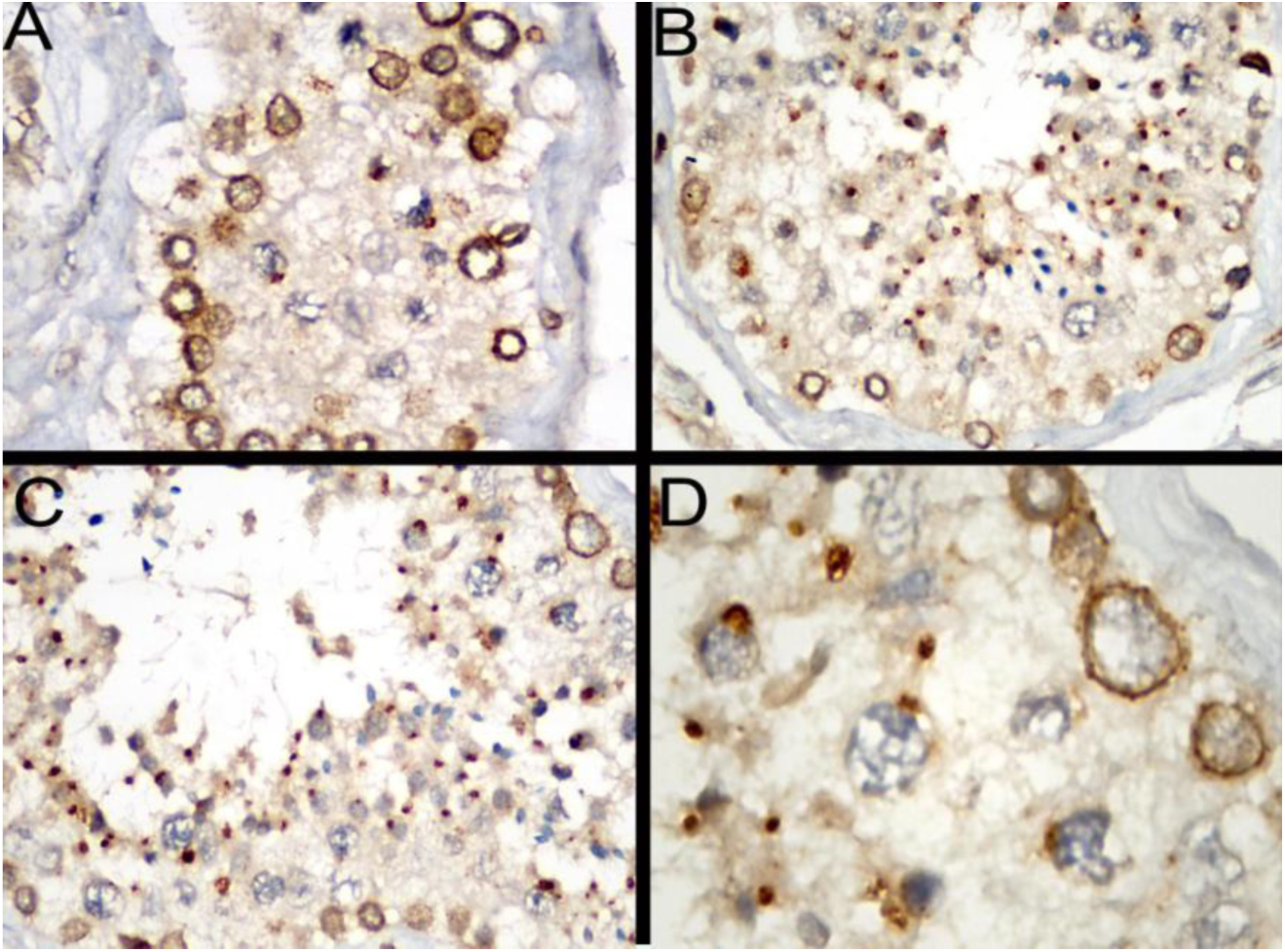
Immunoexpression patterns of DHX37 at different stages of germ cells differentiation. **A.** Perinuclear halo pattern of protein expression in spermatogonia (original magnification ×400). **B.** Adult seminiferous tubule with germ cells showing perinuclear halo in spermatogonia, progressive paranuclear condensation in spermatocytes and spermatids and absence of DHX37 expression in spermatozoa (original magnification ×200). **C.** Numerous spermatids with punctate perinuclear expression and absence of expression in spermatozoa (original magnification ×200) **D.** Magnified images of spermatogonia with perinuclear halo, spermatocyte with paranuclear condensation and spermatids with punctate perinuclear pattern (original magnification ×1000).

## Discussion

The present study analyzed a large cohort of 46,XY DSD patients without defined molecular diagnosis, most of them with gonadal dysgenesis phenotype (63%). Pathogenic allelic variants in the *DHX37* was identified in 16 affected members of 11 families. Deleterious variants are recurrent in familial and sporadic cases of 46,XY GD patients from different ethnicities.

Three variants identified in the present study are located in two highly conserved domains, the Helicase ATP-binding, which is a region associated with RNA-binding activity of the RNA helicases, and the Helicase superfamily c-terminal domain, associated with ATPase activity ^9^. Rare and predicted deleterious *DHX37* variants are more frequently identified in our DSD cohort (14%) than in the public databases, emphasizing that the association of *DHX37* with 46,XY gonadal dysgenesis did not occur by chance (p<0.01).

Since the discovery of *SRY* variants in patients with gonadal dysgenesis in 1990, several genes have been associated with molecular etiology of this disorder^7^. The nuclear receptor subfamily 5 group A member 1 (*NR5A1*) and *MAP3K1* variants are the most frequent causes of 46,XY gonadal dysgenesis ^10–12^.

In this study we found an expressive frequency (10%) of likely-pathogenic variants in *DHX37* in patients with 46,XY gonadal dysgenesis; considering only the ETRS phenotype (micropenis and absence of uni or bilateral testicular tissue) the frequency augmented to 50%.

The potential involvement of *DHX37* in the etiology of male gonadal development defects may be supposed by reports of patients with 12q24.31-33 deletions with syndromic features and genital abnormalities ^13,14^.

*DHX37* encodes an RNA helicase, which is involved in RNA-related process, including transcription, splicing, ribosome biogenesis^15,16^, translation and degradation^17^. Mutations in DEAH Box RNA helicases members have been related to human neurogenic disorders. Disease-causing variants in the DEAH Box Polypeptide 30 (DHX30 [MIM:616423]) gene, were previously described in syndromic patients with global developmental delay (GDD [MIM:166260]), intellectual disability (ID [MIM:614132]), severe speech impairment and gait abnormalities. Functional studies of the altered DHX30 proteins, derived from conserved residue substitutions in the helicase nucleus domain, demonstrated that they affect protein folding or stability interfering with the RNA binding (mutations located in motif Ia) or with ATPase activity (mutations located in motif II and VI)^9^.

Homozygous *DHX37* mutations were previously identified by WES in affected members of two consanguineous families with neurogenetic disorders (developmental delay or intellectual disability) and brain malformations (severe microcephaly or cortical atrophy)^18^. Description of external genitalia or gonadal function of the patients with *DHX37* mutations and neurogenetic disorders was not available.

Neurogenetic disorder was noticed in only one Brazilian ETRS patient, in which the homozygous p.Arg151Trp variant was identified. She was born to a consanguineous Brazilian couple and at age of 4.25 years, she had developmental delay, poor speech and short stature. Unfortunately, her treatment was discontinued after surgery and no information about her was available. According to ACMG guidelines criteria, the p.Arg151Trp variant was classified as a variant of uncertain significance (VUS) (Table S4). This variant is located outside the helicase core region and it was found in heterozygous state in low frequency (0.02%) in the 1000 Genomes Database. Although these facts suggest that the heterozygous p.Arg151Trp variant has no significant phenotypic effect, we speculate a possible pathogenic role when it is in homozygous state, similarly to previous described for mild homozygous *NR5A1* mutations in 46,XY DSD patients ^19,20^.

The unique p.Thr304Met variant and the recurrent p.Arg308Gln variant were located in a region associated with RNA-binding activity of the RNA helicases^9^. The p.Arg308Gln variant was identified in 8 affected members of six families with ETRS but a founder effect for this variant was ruled out. Two patterns of inheritance of the p.Arg308Gln variant were revealed by segregation analysis: the sex-limited autosomal dominant mode found in 4 families and autosomal dominant with incomplete penetrance found in the Family 1. In the literature, different inheritance patterns had already been identified in 46,XY gonadal dysgenesis kindreds^21^, including the description of asymptomatic male members carriers of pathogenic variants of genes involved in the testicular determination as *SRY* and *NR5A1*^19,22,23^

The heterozygous p.Arg674Trp, identified in six affected members of three unrelated families with ETRS and partial GD phenotype, was transmitted as a sex-limited autosomal dominant mode. The Residue Arg674 is located in the Motif VI, other highly conserved position of the RNA helicases of Helicase associated with ATPase activity^9^.

DEAD-box proteins have been associated to the control cellular growth, proliferation and differentiation, and are known to be implicated in embryogenesis and spermatogenesis^17,24^. DExD/H-box RNA helicases regulate reproductive–related genes by modulating mRNA structures^25^. RNA expression profiles of *DHX37* in human testicular cancer cells are higher than in other tissue (The Human Protein Atlas – Pathology), suggesting that the DHX37 may be involved in the regulatory process of the testis cells proliferation.

Expression analysis of RNA helicase genes in channel catfish gonads demonstrated a sex-specific expression pattern. The *DEAD-box RNA helicase 5 (DDX5/p68* [MIM:180630]), *DEAD-box RNA helicase 21 (DDX21* [MIM:606357]) and *DEAH box Polypeptide 9 (DHX9* [MIM:603115]) were highly expressed during the critical period of male catfish gonadal differentiation indicating a role in male gonadal development^26^.

In humans, the DHX37 protein is express in adults ovarian stroma and seminiferous duct cells (The Human Protein Atlas Database). In our study, the immunohistochemistry analysis of normal testicular tissue from stillborn, pubertal and adult males revealed that DHX37 was expressed during different stages of germ cell maturation. These findings suggest that DHX37 protein might have a role in the germ cell development. Although it is still unclear how germ cells contribute to the development and function of somatic cells in the testis, there is an elaborate paracrine cell-cell network transporting signaling molecules between germ cells and Sertoli cells^27^. Indeed, *in vitro* studies have shown that there is a bidirectional trafficking between Sertoli and germ cells, and that each cell type regulates the function of the other^28–31^.

Based on protein interaction networks we hypothesize that the role of DHX37 in male testicular development involves the regulation of the serine/threonine protein kinase encoded by *V-Akt Murine Thymoma Viral Oncogene Homolog 1 (AKT1 [MIM: 164730])*. Our analysis suggests that DHX37 may interact with DEAH-box RNA helicase 15 (DHX15 [MIM: 603403]) and DDX5/p68, known regulators of *AKT1*^32,33^. This serine-threonine kinase is expressed in germ cells and Sertoli cells and plays an important role in the proliferation, differentiation and maintenance of testicular tissue, including regulation of apoptosis in male germ cells^34^. In addition, AKT1 also regulates the Mitogen-Activated Kinase 1 (MAP3K1)-RHOA/ROCK complex and, thus, the phosphorylation of p38/Mitogen-Activated Protein Kinase 14 (MAPK14 [MIM: 600289]), ERKs (Mitogen-Activated Protein Kinase, MAPK3 [MIM: 601795]; MAPK1 [MIM: 176948]), which are essential for SOX9 (MIM:608160) activation and testis determination (Figure 4)^35–39^. Further studies are necessary to clarify the precise role DHX37 plays in the signaling network that regulates testicular development and maintenance.

**Figure 4.**
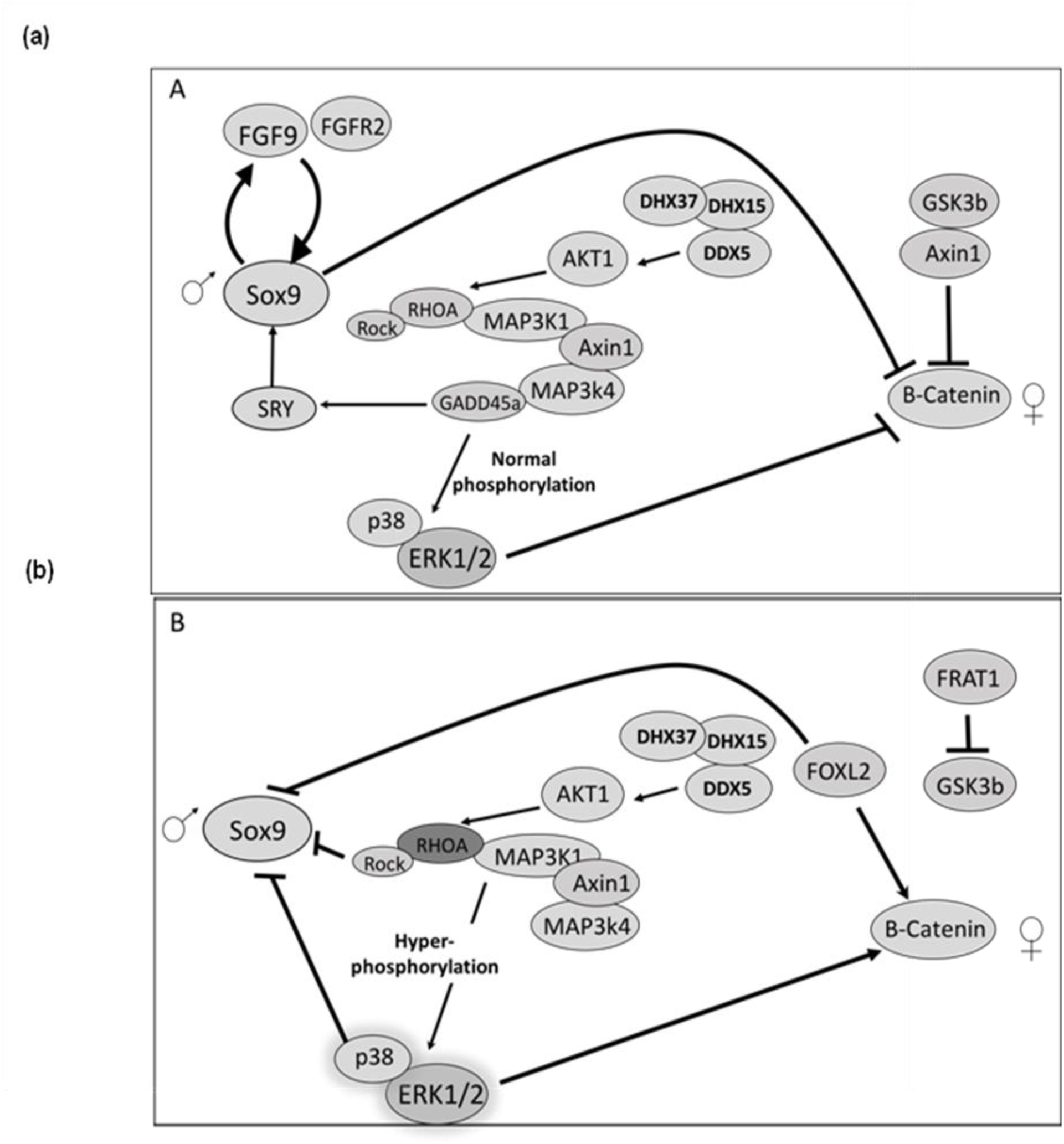
Representation of proposed interactions of DHX37 in the male gonadal development pathway. **A.** Normal testicular development - Interaction of DHX37 with DHX15 and DDX5 helicases leading to normal phosphorylation of AKT1, which interacts with the RHOA/ROCK-MAPK (MAP3K1/AXIN1/MAP3K4/GADD45a) pathways, as well as of ERK1/2 and p38 MAPK protein kinases. ERK1/2 and p38 kinases block the beta-catenin pathway and promote SOX9 stimulation leading to testicular development. **B.** Abnormal testicular development - Changes in DHX37 could interfere with the normal interaction between helicases and cause decreased AKT1 phosphorylation. In this condition, the RHOA/ROCK-MAPK signaling was increased and ERK1/2 and p38 hyperphosphorylation leads to SOX9 repression through beta-catenin/FOXL2 activation, resulting in abnormal testicular development. Factors that promote the action of targets are shown as black arrows. Factors that disrupt the action of targets are shown as broken lines.

## Supporting information

## Acknowledgments

This work was supported by grants from the Conselho Nacional de Desenvolvimento Científico e Tecnológico (CNPq, Grant No. 305743/2011-2), the Fundação de Amparo à Pesquisa do Estado de São Paulo (FAPESP, Grants No. 05/04726-0, 07/512156, 10/51102-0, 2013/02162-8 and 2014/50137-5), and by the institutional fellowship grant from the Coordenação de Aperfeiçoamento de Pessoal de Nível Superior (CAPES/ PNPD). The authors are very grateful to Dr. Frederico Moraes Ferreira for his technical assistance with the *in silico* prediction analysis, and Dr. Beverly M. Yashar and Dr. John Park for medical care of patient F5:II-1.

## Web Resources

1000 Genomes Project, http://www.internationalgenome.org/

ABraOM (http://abraom.ib.usp.br/search.php)

ANNOVAR (http://annovar.openbioinformatics.org/en/latest/)

biobambam2 suite (https://launchpad.net/biobambam2)

Beagle software (http://faculty.washington.edu/browning/beagle/beagle.html)

Biogrid (https://thebiogrid.org/)

CADD (http://cadd.gs.washington.edu)

dbSNP (https://www.ncbi.nlm.nih.gov/SNP/)

ESP6500 (http://evs.gs.washington.edu/EVS/)

ExAC Browser (http://exac.broadinstitute.org/)

FATHMM (http://fathmm.biocompute.org.uk)

FastQC, http://www.bioinformatics.babraham.ac.uk/projects/fastqc/

Freebayes (https://github.com/ekg/freebayes)

GATK, https://software.broadinstitute.org/gatk/

GenBank, https://www.ncbi.nlm.nih.gov/genbank/

GERP++ (http://mendel.stanford.edu/SidowLab/downloads/gerp/)

Human Protein Atlas (https://www.proteinatlas.org/)

IGV (http://www.broadinstitute.org/igv/)

Mutation Taster (http://www.mutationtaster.org)

Mutation Acessor (http://mutationassessor.org/r3/)

OMIM, https://www.omim.org/

PhyloP (http://compgen.cshl.edu/phast/background.php)

PolyPhen2 (http://genetics.bwh.harvard.edu/pph2/)

SIFT (http://sift.jcvi.org)

SiPhy (http://portals.broadinstitute.org/genome_bio/siphy/index.html)

String (https://string-db.org/)

