## Supplementary material for "DEAH-box helicase 37 (*DHX37*) defects are a novel molecular etiology of 46,XY gonadal dysgenesis spectrum"

**Supplementary data**

Details of the case reports, exome capture, sequencing and variant filtering and immunohistochemical expression analysis are available in the Supplemental Data that includes four tables.

**Patients**

**Family 1.** A Brazilian family with two male siblings born to non-consanguineous parents presenting micropenis and bilateral cryptorchidism (Table 1). No family history of atypical genitalia or infertility was reported. Both pregnancies and births were uneventful. We evaluated the eldest affected child (F1:III-2; Fig. 1) when he was 2.18 years old. Physical examination showed a micropenis (penile length of 1.5 cm, -4.0 SD), empty scrotum and nonpalpable gonads. After hCG stimulation test, no increase in prepubertal testosterone or androgen precursor levels was observed (Table 1). MRI did not reveal Müllerian structures. During exploratory laparoscopy, no uterus was identified, and the bilateral streak tissue was resected. The histopathological study revealed a small bilateral area of dysgenetic gonadal tissue and the presence of bilateral epididymis and Müllerian ducts that would differentiate into fallopian tubes (Table 1). His younger brother (Subject F1:III-3; Fig. 1) was first evaluated at 1.8 years of age due to primary features of a small penis (Table 1). His birth was normal. Genital examination showed a micropenis (penile length of 2.0 cm; -3.4, SD), glans-urethral meatus and nonpalpable gonads. MRI did not identify Müllerian structures. During laparoscopic surgery, no uterus was identified and small gonadal tissues, located in the internal inguinal rings, were resected. The histological study identified fibrous connective tissue with scattered degenerate seminiferous tubules showing hydropic degeneration of

Sertoli cells on the right side and no gonadal tissue on the left side. Epididymis, ductus deferens and rete testis were present bilaterally (Table 1).

**Family 2.** The proband of Family 2 (Subject F2:II-4; Fig. 1), a Brazilian family, was born to non-consanguineous parents and first evaluated at the age of 14 years due to the presence of a micropenis. Information about pregnancy and delivery was not available. External genitalia examination showed a micropenis (penile length of 1.5 cm; -7.0 SD), glans-urethral meatus and nonpalpable gonads. No increase in prepubertal testosterone levels was observed after hCG stimulation test. Exploratory laparoscopy revealed a normal epididymis, but no uterus or gonadal tissue. The histological study revealed presence of epididymis, ductus deferens, rudimentary Müllerian ducts and absence of gonadal tissue (Table 1). His niece was also born with micropenis (penile length of 1.0 cm, -4.6 SD), glans-urethral meatus and nonpalpable gonads, but was reassigned and raised as a female (Subject F2:III-1; Fig. 1). During exploratory laparotomy, atrophic gonadal tissue on the right side was resected and no uterus was visualized. The histological study identified presence of fibrous connective tissue with a very small area of dysgenetic gonadal tissue on the right side and no gonadal tissue on the left side. A normal epididymis, ductus deferens and rudimentary Müllerian ducts were identified (Table 1). No consanguinity or family history of infertility was reported.

**Family 3.** Two male siblings of a Chilean family were evaluated. They were born to non-consanguineous parents (Table 1) and presented the primary features of small penis. The parents reported no family history of atypical genitalia or infertility. Both pregnancies and births were uneventful. The eldest affected child (F3:II-1; Fig. 1) was evaluated at 6 months of age. Physical examination showed presence of a micropenis (penile length of 0.5 cm, -4.2 SD), an empty scrotum and nonpalpable gonads. During exploratory laparoscopy, no uterus was identified and the bilateral streak tissue was resected. The histopathological study revealed a small bilateral area of dysgenetic gonad, bilateral epididymis and absence of

Fallopian tubes (Table 1). His younger brother (F3:II-2; Fig. 1), evaluated at age of 24 days, presented a similar phenotype. His physical examination revealed presence of a micropenis (penile length of 0.8 cm; -3.9 SD), an empty scrotum and nonpalpable gonads. During laparoscopic surgery, an uterus was revealed and the bilateral streak tissue was resected. The histopathological study revealed a small bilateral area of dysgenetic gonad consisting of a tubular structure similar to the one of a Fallopian tube (Table 1).

**Family 4.** Two male siblings and one maternal uncle with a small penis (Table 1) of an Argentine family were evaluated. All pregnancies and births were uneventful. The eldest affected child (F4:III-1; Fig. 1) was first noticed to have a micropenis (penile length of 1.2 cm, -5.7 SD), distal hypospadias, a hypoplastic, empty scrotum and nonpalpable gonads shortly after birth. Exploratory laparoscopy did not reveal presence of an uterus or gonads. The histopathological study revealed absence of testicular tissue, bilateral epididymis and rudimentary Fallopian tubes (Table 1). A similar phenotype was observed in his younger brother (F4:III-2; Fig. 1), who was 4 years of age at the time of evaluation. The physical examination revealed presence of a micropenis (penile length of 2.5 cm; -3.3 SD) and nonpalpable gonads. During laparoscopic surgery, an uterus was not identified and the bilateral streak tissue was resected. The histopathological study revealed presence of epididymis and rudimentary Fallopian tubes (Table 1). The maternal uncle (F4:II-4; Fig. 1) was evaluated at 2 years of age. The physical examination revealed presence of a microphallus (penile length of 1.5 cm; -4.0 SD), distal hypospadias, left testis in the scrotum and a palpable structure in the right inguinal region. Laparoscopic surgery was performed at 15 years of age, but did not reveal the presence of an uterus. Gonadal tissue was not identified on the right side, and the histopathological study of the left testicular biopsy samples revealed testis with a thick tunica albuginea and a small area of testicular parenchyma that presented tubules with peripheral hyalinization and undifferentiated intratubular germinal neoplasia; epididymis and deferent duct were identified on the right side (Table 1).

**Sporadic case F5:II-1.** (Fig. 1) The proband was the first child born from a non-consanguineous Chinese couple. He was first noticed to have an atypical genitalia shortly after birth. Ultrasonography showed no evidence of scrotal or pelvic gonadal tissue. A lobulated structure was observed between the bladder and rectum, possibly similar to a Müllerian remnant. Genetic testing revealed a normal 46,XY chromosomal microarray. Following an hCG stimulation test, presented no significant increase in testosterone or hormonal precursor levels. At the age of 3 years, an external genitalia examination showed a phallic structure (penile length of 1.5 cm, -4.0 SD), subcoronal hypospadias and nonpalpable gonads in the scrotum or inguinal areas. Laparoscopy identified a poorly formed epididymis bilaterally and small, dysplastic appearing gonads, which were removed. The pathological evaluation identified atrophic epididymal tissue and vas deferens, no evidence of Müllerian structures and no gonadal tissue (Table 2). Family history was negative for atypical genitalia or infertility.

**Sporadic case F6:II-1.** (Fig. 1) The Brazilian patient underwent treatment at 30 years of age (Table 2). She was born with micropenis and was assigned to the male gender, but since childhood she has identified herself as a female. Information about pregnancy and delivery was not available. The patient denied the development of secondary sexual characteristics. Sex reassignment (male to female) was carried out after psychological evaluation. Physical examination revealed the following features: Height- 171 cm, BMI- 22.4 $kg/cm^2$, breast-Tanner stage I, pubic hair- Tanner stage III, external genitalia with a micropenis (penile length of 3.5 cm, -6.0 SD), glans-urethral meatus and nonpalpable gonads. No increase in testosterone or androgen precursor levels was observed after hCG stimulation test. Pelvic ultrasonography did not identify the presence of an uterus and gonads. A feminizing genitoplasty was performed. During exploratory laparoscopy no uterus was found and streak gonadal tissue was removed. The histological analysis identified no gonadal tissue. Fallopian tubes and epididymis were identified bilaterally (Table 2).

**Sporadic case F7:II-1.** (Fig. 1) The patient was born to a consanguineous Brazilian couple. The parents were first cousins, and the family history revealed that their son had speech impairment. The child was reassigned and raised as a female. The child was first evaluated at 4.25 years of age due to atypical genitalia. The pregnancy and delivery were normal. She presented with signs of malnutrition and growth retardation, developmental delay and poor speech. The external genitalia showed a micropenis (penile length of 0.6 cm, -5.6 SD), glans-urethral meatus and nonpalpable gonads. No increase in prepubertal testosterone or androgen precursor levels was observed after hCG stimulation test. Pelvic ultrasonography did not reveal the presence of an uterus and gonads. During exploratory laparotomy, an uterus was not revealed, and streak gonadal tissue was removed. The histological analysis revealed no gonadal tissue on the right side and a small nest of dysgenetic gonad on the left side. Epididymis, ductus deferens and Fallopian tubes were identified bilaterally (Table 2).

**Sporadic case F8:II-1.** This Brazilian child was referred at 7.7 years old to the Hospital das Clinicas due to short vagina and absence of uterus. She is the only child of a non-consanguineous couple and was born after an uneventful pregnancy. A 46,XY karyotype was identified. Laparotomy was performed and no Mullerian derivatives were identified, and the streak gonadal tissue was removed. There were no similar cases in her family.

**Sporadic case F9:II-1**. (Fig. 1) This Brazilian patient was referred to our clinic at the age of 35 years. She was born with atypical genitalia and underwent gonadectomy and feminizing genitoplasty at 2 years of age at another hospital (Table 2).

**Sporadic case F10:II-1**. (Fig. 1) The Brazilian patient was referred to our clinic at the age of 19 years old (Table 2). She was born with a micropenis and was assigned male at birth.

However, since childhood she had a female gender identity. Sex reassignment (male to female) was carried out after psychological evaluation. Physical examination revealed a micropenis (penile length of 5.5 cm, -4.9 SD), glans-urethral meatus and nonpalpable gonads. No increase in testosterone or androgen precursor levels was observed after hCG stimulation test. Pelvic ultrasonography did not identify the presence of uterus and gonads. A feminizing genitoplasty was performed. During exploratory laparoscopy no uterus was found and streak gonadal tissue was removed. The histological analysis identified no gonadal tissue (Table 2).

**Sporadic case F11:II-1.** This Brazilian child was referred to our Hospital at 3.7 years of age because of an atypical genitalia characterized by a single perineal opening and bilateral cryptorchidism. An increase in prepubertal testosterone (337 ng/dL) without an increase in other androgen precursor levels was observed after hCG stimulation test. Laparoscopy was performed and dysgenetic gonadas associated with fallopian tubes were identified.

#### Methods

*Whole-exome sequencing:* For sequence alignment, indexing of the reference human genome (GRCh37.p11; UCSC Genome Browser hg19), variant calling, and annotation, a pipeline based on Burrows-Wheeler Aligner (BWA-MEM) v0.7.1(1), Genome Analysis Toolkit (GATK) v3.8 software (2) package, and ANNOVAR (3) was used, respectively. In order to specifically select disease-causing variants, we excluded those with minor allele frequency (> 0.1%) in publicly available population databases, such as ExAC (4), 1000 Genomes (5), ESP6500 (6), dbSNP (7), and AbraoOM (8)-Brazilian Genomic Variants, and in an in-house massively parallel sequencing database of 600 Brazilian patients with no DSD (SELA). To assess the possible impact on the protein structure and function of the novel non-synonymous variants we used several *in silico* algorithms (SIFT(9), PolyPhen2(10), Mutation Taster (11), Mutation Assessor (12), FATHMM (13), LRT (14), LR score (15), and radial SVM (16)) and by the conservation scores (GERP++ (17), SiPhy (18), PhyloP (19), CADD score(20)).

*Target massively parallel sequencing (TMPS):* Their entire coding regions and respective 50-bp boundaries were captured using custom Sure Select Target Enrichment System Kit (Agilent). Sequencing was performed on the Illumina MiSeq platform. Paired-end reads (2 x 300) were aligned to the hg19 assembly of the human genome with BWA-MEM. Aligned reads were sorted and converted into BAM format using the bamsort tool from the biobambam2 suite (21). Mean coverage was over 95× for all samples, and more than 96% of the RefSeq gene coding regions were covered at 20× or deeper. Single-nucleotide variants and small INDELs were simultaneously called in all samples using Freebayes (22). Variant annotation was performed using ANNOVAR.

15. Dong C, Wei P, Jian X, et al. Comparison and integration of deleteriousness prediction

methods for nonsynonymous SNVs in whole exome sequencing studies. *Hum Mol Genet*. 2015;24(8):2125-2137. doi:10.1093/hmg/ddu733.

#### Supplementary Tables

**Table S1.** Genes associated with DSD studied by target massively parallel sequencing

| | | | | | | |
|---|---|---|---|---|---|---|
| **Genes involved in the gonadal determination** | | | | | | |
| *AMH* | *DMRT2* | *LHX1* | *FOXL2* | *SOX9* | *PBX1* | *WWOX* |
| *CBX2* | *FGF9* | *MAMLD1* | *NANOS3* | *SRY* | *AXIN1* | *WT1* |
| *CITED2* | *FGFR2* | *MAP3K1* | *PTDGS* | *STAG3* | *PAX2* | *LHX9* |
| *DHH* | *GATA4* | *NR0B1* | *RSPO1* | *WNT4* | *NANOS2* | *PAPPA* |
| ***DHX37***# | *GSK3B* | *NR5A1* | *RSPO2* | *WXT1* | *TCF21* | *STRA8* |
| *DMRT1* | *GDF9* | *BMP15* | *SOX3* | *ZFPM2* | *TES* | |
| **Genes associated with hormonal synthesis** | | | | | | |
| *AKR1C2* | *AKR1C3* | *AKR1C4* | *CYP17A1* | *CYP19A1* | *HSD11B1* | *HSD17B3* |
| *HSD3B2* | *POR* | *SRD5A2* | *STAR* | *RAC1* | *CYP21A2* | |
| *Genes associated with hormonal receptors* | | | | | | |
| *AMHR2* | *ESR2* | *FSHR* | *LHCGR* | *ESR1* | *AR* | |
| *Genes associated with syndromic genital malformations* | | | | | | |
| *ARX* | *ATRX* | *CDH7* | *HNF1B* | *CDH7* | | |

### DHX37 was not previously associated with DSD. It has included in our the targeted sequencing panel after had been identified in two Brazilian individuals with familial ETRS

**Table S2.** *In silico* prediction analysis of DHX37 allelic variants identified in 46,XY DSD patients

| Subjects | Nucleotide changed | AA changed | Functional domain | *In silico* prediction tools | | | | | |
|---|---|---|---|---|---|---|---|---|---|
| | | | | Mutation Taster | Mutation Assessor | SIFT | Polyphen-2 | CADD | GERP |
| F1, F2, F5, F6, F8 | c.923C>T | p.Arg308Gln | Helicase ATP-binding | Pathogenic (score: 0.999) | High functional impact (score: 4.38) | Protein function affected (score 0.001) | Probably deleterious (score 1.000) | 35 | 5.3 |
| F3,F4,F10, F11 | c.2020G>A | p.Arg674Trp | Helicase superfamily C-terminal domain | Pathogenic (score 1.000) | Middle functional impact (score: 4.835) | Protein function affected (score 0.001) | Disease cause (score 1.000) | 33 | 4.2 |
| F7 | c.451G>A | p.Arg151Trp | unknown | Pathogenic (score: 0.999) | Middle functional impact (score: 2.85) | Protein function affected (score 0.001) | Possible deleterious (score 0.669) | NA | NA |
| F9 | c.911C>T | p.Thr304Met | Helicase ATP-binding | Pathogenic (score 1.000) | High functional impact (score: 4.45) | Protein function affected (score 0.001) | Disease cause (score 1.000) | 29.8 | 5.3 |

**Table S3.** *DHX37* missense allelic variants identified in 46,XY DSD patients and their frequency in population databases

| **Families** | **cDNA position** | **AA change** | **Phylogenetic Conservation*** | **State** | **dbSNP** | **MAFs in population databases** | | | | |
|---|---|---|---|---|---|---|---|---|---|---|
| | | | | | | **1000 Genomes** | **ExAC** | **gnomAD** | **ABraOm** | **ESP6500** |
| F1,F2, F5, F6, F8 | c.923 C>T | p.Arg308Gln | Highly conserved | Heterozygosis | Not available | Absent | Absent | 0.00006677 Non-Finnish European | Absent | Absent |
| F3, F4, F10, F11 | c.2020G>A | p.Arg674Trp | Highly conserved | Heterozygosis | Not available | Absent | Absent | Absent | Absent | Absent |
| F7 | c.451G>A | p.Arg151Trp | Middle conserved | Homozygosis | rs577400960 | 0.00019972 (1/5007) South Asian | 0.00012703 (3/23616) Non-Finnish European and South Asian | Absent | Absent | Absent |
| F9 | c.911C>T | p.Thr304Met | Highly conserved | Heterozygosis | Not available | Absent | Absent | Absent | Absent | Absent |

**Table S4.** Pathogenicity classification of *DHX37* variants according to the ACMG guidelines

| Families | Nucleotide changed | AA changed | Population Data | Computational and prediction data | De novo data | Other data | Classification |
|---|---|---|---|---|---|---|---|
| F1,F2, F5, F6 | c.923C>T | p.Arg308Gln | PM2 [a] | PP3 [b] | PS2 [c] | PM1[d] PP4[e] | Likely Pathogenic |
| F3, F4, F10,F11 | c.2020G>A | p.Arg674Trp | PM2 [a] | PP3 [b] | | PM1[d] PP4[e] | Likely Pathogenic |
| F7 | c.451G>A | p.Arg151Trp | PM2 [a] | PP3 [b] | | | VUS |
| F8 | c.911C>T | p.Thr304Met | PM2 [a] | PP3 [b] | | | VUS |

PM2: moderate piece of evidence for pathogenicity; PP3: supporting evidence for pathogenicity by computational (in silico) data;
PS2: strong support for pathogenicity when the de novo variants; PP4: supporting evidence using phenotype; PM1: pathogenic moderate;
VUS: Variant of Uncertain Significance.

a Absent from controls (or at extremely low frequency if recessive) in Exome Sequencing Project, 1000 Genomes Project, or Exome Aggregation Consortium.

b Multiple lines of computational evidence support a deleterious effect on the gene or gene product (conservation, evolutionary, splicing impact, etc.)

c De novo (both maternity and paternity confirmed) in a patient with the disease and no family history.

d Located in a mutational hot spot and/or critical and well-established functional domain (e.i., interaction with RNA) without benign variation.

e Patient's phenotype or family history is highly specific for a disease with a single genetic etiology.
